# Is the distribution of the Acorn Woodpecker (*Melanerpes formicivorus flavigula*) associated with oaks and granaries? A local study in an urban area in northern South America

**DOI:** 10.1101/765487

**Authors:** Hana Londoño Oikawa, Paulo C. Pulgarín-R

## Abstract

Abiotic and biotic factors are known to be key in limiting the geographical distribution of species. However, our understanding on the influence of habitat heterogeneity on ecological interactions and behavior in tropical animals is limited. We studied groups of Acorn Woodpeckers (*Melanerpes formicivorus flavigula*) in urban and rural areas in northern South America to understand how habitat and resource requirements (food storage structures) influences patterns of distribution across the Aburrá Valley, in the northern area of the Central Andes of Colombia. Using focal observations of 10 different groups over nearly a two-year period, we estimated territory size, habitat use, and described the use and presence of granaries. We found that territory size, tree diversity, and the use of granaries varied among groups. Accordingly, Acorn Woodpeckers use a wide variety of tree species to make cavities, to feed and to build granaries for social interactions. Our study supports the hypothesis that Acorn Woodpeckers do not rely on the Colombian Oak (*Quercus humboldtii* Bonpl.) for feeding, nesting or foraging in the Aburrá Valley, and that the construction of granaries to store food is present in urban populations, despite the lack of strong seasonal changes in tropical areas. We suggest that the distribution of the Acorn Woodpecker in our study area is strongly associated with one particular species of tree, *Albizia carbonaria* Britton, and the behavior of granaries construction might be hardwired in this species for the maintenance and cohesion of family groups.

## Introduction

The geographic distribution of different groups of animals and plants are determined by abiotic (e.g. precipitation and temperature) and biotic (e.g. competition, parasitism or mutualism) factors (Sexton et al. 2009, Schmitt et al. 2012). How variation in such factors contribute to explain range limits in different organisms are major subjects of study in biogeography, ecology and conservation, particularly when we live in a world where natural environments are rapidly being transformed by humans (Quammen 1997). Likewise, behavioral, ecological, and life history traits usually vary in species that are widely distributed (Jetz & Rubenstein 2011), suggesting that organisms respond and adjust to variation of habitat, climate and interactions with other taxa (Rubenstein & Lovette 2007). In animals and particularly in birds, there have been many studies examining whether ecological characteristics, or climatic variables are the key limiting factor governing the distribution of species (Araújo & Luoto 2007, Freeman & Mason 2015). Species such as the American Krestel (*Falco peregrinus*), Rufous-collared Sparrow (*Zonotrichia capensis)*, Great Tit (*Parus major*) and the Andean Condor (*Vultur gryphus*) are examples of bird species with wide distributional ranges encompassing different climates and habitats; these and other taxa are experiencing ecological, behavioral, and evolutionary changes in environments created by human activities, including urbanization (Wallace & Temple 1987, Bonier 2012, Lougheed et al. 2013, Bosse et al. 2017)

In urbanized environments in the tropics, particularly in northern South America, there is a lack of studies on bird species that are widespread and that are able to use many types of habitats, or to persist in environments highly modified by anthropogenic activities (Martin & Bonier 2018). What allows certain species to remain and other to disappear is still unknown. For instance, some species might be better competitors than others or are more generalist in their diets. Although there are local studies on native and introduced species in urban environments, such as foraging behavior (Delgado et al. 2005, Delgado 2007, Delgado & Calderón 2007, Osorio & Marín 2016); nesting biology (Sedano et al. 2008); interactions with other species (Garcés et al. 2012); or abundance and reproduction (Sánchez et al. 2016), there are a lack of studies examining how species who are widely distributed have modified their behavior and use of habitat to thrive in local conditions (Boyce & Martin 2017).

The Acorn Woodpecker (*Melanerpes formicivorus*, Aves: Picidae) has a long and discontinuous distribution that ranges from the western North America to northern South America in Colombia, where it occupies montane-forest in the three Andean Cordilleras (Winkler & Christie 2019). Extensive studies about its diet (Stacey 1981, Rosas et al. 2008), parental care (Koenig & Walters 2011), population ecology (Koenig & Mumme 1987) and use of habitat (Stacey 1979, Ligon & Stacey 1996) have been done in North and Central America. A well known aspect is their strong relationship with different oak species of the genus *Quercus* and *Notholithocarpus;* from which they rely on as a food source (Scofield et al. 2011) and as nest cavity sites (Koenig & Walters 2014), allowing them to survive during the winter season (Koenig & Haydock 1999). A behavior associated with the use of oaks is the accumulation of acorns or seeds in small holes in the bark of these trees (MacRoberts & MacRoberts 1976). These storage trees or structures are called granaries, and are considered one of the key factors for the social and population structure of this species ((Koenig & Walters 2014).

In Colombia, where the subspecies *Melanerpes formicivorus flavigula* (one of seven subspecies across the whole range) occupies the southern limit of its distribution (Hilty & Brown 1986), there are a handful of studies examining their diet and distribution at local and regional scales. Such studies show: (1) descriptions of cavities and feeding behavior of a population in southern Colombia (Miller 1963); (2) the interaction with the Colombian Oak (*Quercus humboldtii* Bonpl.) and the Buff-tailed Coronet (*Boissonneaua flavescens*) (Kattan & Murcia 1985); (3) the food habits and social organization (Kattan 1988); and (4) that according to a species’ distribution model, the Acorn Woodpecker is limited by the distribution of Colombian Oak in the Colombian Cordilleras (Freeman & Mason 2015). The prevailing consensus is that *M. formicivorus flavigula* is ecologically associated with Q. *humboldtii*, and that the use and construction of granaries does not seem to be a recurrent behavior in Colombian populations.

Here, we study the local distribution and habitat use of *M. f. flavigula* in the Aburrá Valley region of northern South America (a complex region with natural and urban environments). In particular, we aimed to (1) describe territory sizes and group composition; (2) to identify which species of trees are being used by *M. f. flavigula* for foraging and cavity construction; and (3) to determine if the species is constructing and using granaries. Based on our own observations, we hypothesized that *M. f. flavigula* does not rely and is not limited by the Colombian Oak (*Quercus humboldtii*) at least in the Aburrá Valley, and that the behavior of granary construction is maintained in tropical populations of the Acorn Woodpecker despite apparent lack of strong seasonal changes.

## Methods

### Study Region

Our study area is located in the Tropical Andes, at the north portion of the Central Cordillera of the Colombian Andes (Figure 1). It is situated within a natural region called the Aburrá Valley, which has an extension of ~70 km of length and 25 km of width. The Medellín River runs (south-north) through the valley, across ten municipalities: Caldas, Sabaneta, La Estrella, Envigado, Itagüí, Medellín, Bello, Copacabana, Girardota and Barbosa. The Acorn Woodpecker groups we studied were located in three of those municipalities (Envigado, Medellín and Bello), which have been rapidly urbanized, but are also important areas for endemic and endangered animals and plants biodiversity (Palacio et al. 2006, Delgado & Correa 2013, Sánchez et al. 2014, Varón & Morales 2016). We worked at elevations from 1500 to 2600 m. The mean temperature in the study area is 22.5°C, and the mean precipitation is 1450 millimeters per year. The study area was originally covered with premontane and montane humid forest (Espinal 1992). However, most of the original vegetation has been cleared and urbanized, and only few remnants of the native forest remain.

**Figure 1.**
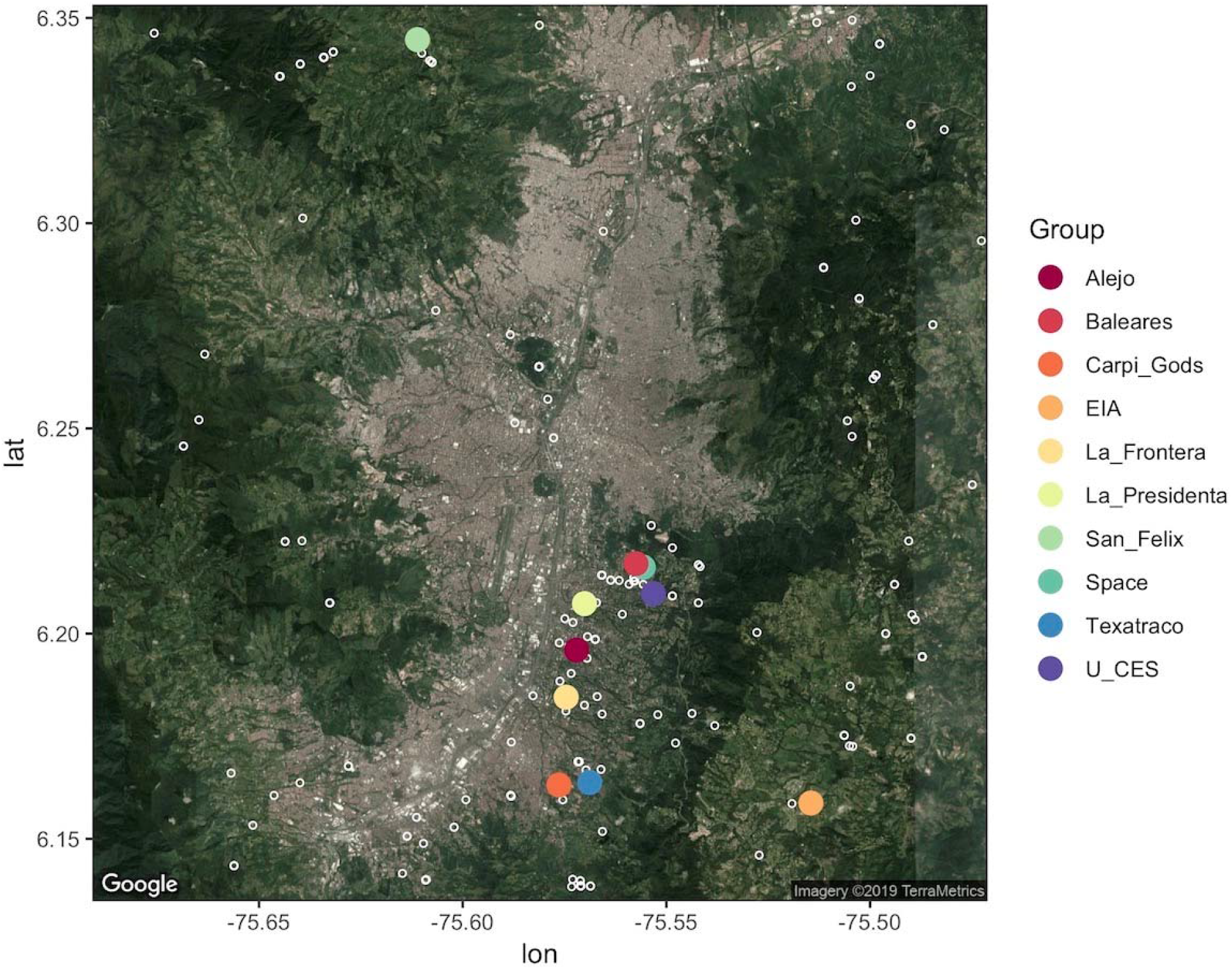
Study area and distribution of focal (n=10) study groups of the Acorn Woodpecker (*Melanerpes formicivorus flavigula*) in the Aburrá Valley and adjacent areas. Open white circles represent eBird records by March 2019.

### Data Collection

#### Group selection

We conducted surveys and gathered data from December 2015 to August 2017. First, we located groups of *M. f. flavigula* by visiting areas where this species had been previously sighted by ourselves or by local ornithologists (e.g. eBird). When we detected one or several individuals, we geo-referenced the location, and expended several hours/days understanding flying paths and core foraging areas. Our total effort time was 293 hours (~ 29.27 ± 11.53 hours/per group). Although our sampling spanned over one and a half years, we did not collect data for several months (i.e. January and April 2016, and May, June, and December 2017). Our initial exploration suggested that in most sites where the species was present, we also detected *Albizia carbonaria* trees (pers. obs); as a consequence, we began looking for and denoting the presence of this tree species in our searching areas. In several cases (five out of ten) we found groups of *M. f. flavigula* associated with this tree species.

#### Group sizes and structure

For each group, we recorded the maximum number of individuals, and whenever possible the sex and age of each individual based on morphological traits through direct observations or with binoculars (MacRoberts & MacRoberts 1976, Short 1982, Hilty & Brown 1986). Although we identified chicks and juveniles (in some territories) by their plumage and their vocalizations inside nest cavities (Winkler & Christie 2019), we didn’t include them in the number of individuals per territory because we did not carry out a rigorous monitoring on them. Nonetheless, we included this information in our overall results, since many aspects of the reproduction and behavior of the Acorn Woodpecker remains to be studied in Colombian populations (Miller 1963, Hilty & Brown 1986, Kattan 1988, Koenig & Walters 2014).

We dedicated between three to seven days (non-consecutive) in each group to gather data to estimate territory size, group size and to collect information about foraging, habitat and behavior. To get a conservative estimation on the number of individuals in each site/territory, we counted the number of individuals present at the same time at their main foraging tree (granary or cavity tree, which often was the most visited place within a group site) early in the morning (06:00 to 8:00 h), when most individuals were coming out of the cavities or were vocally active. Observations were made using Vixen ATREK 10 × 25, and Nikon Monarch 3 10 × 42 binoculars, by one or two simultaneous observers. To avoid affecting the birds behavior we tried to hide or take special precautions, but since these populations seem to be accustomed to human presence, we regularly got close to individuals and trees in order to get a better understanding of their behavior, diet or other information. In some cases, where two territories were in close proximity (1 to 2 km), we collected data simultaneously in both groups to avoid overestimating territories, making sure each group was independent. As banding birds was not included in the methods used, we took special precaution for observing groups having certainty of them being independent from each other.

#### Tree diversity

To characterize the habitat used by *M. f. flavigula*, we identified species of trees where birds were seen nesting, foraging, perching and feeding. We also quantified the number of individual trees frequently used in each group. Additionally, we documented if the birds were using trees for feeding (insects, fruits, seeds), as cavities, or as granaries. Also, we classified whether the tree species were native or exotic with the help of a local botanist and identification field guides (Callejas 2011, Varón & Morales 2016).

#### Use of granaries and feeding preferences

In each territory, we counted the number of granaries, and whenever possible documented the activity around these structures. We recorded if the granary was active (e.g. active for food store or to get food), and if the tree used as granary was dead or alive. Whenever possible, we characterized and quantified the food preferences in each group by counting opportunistically the number of times we observed individuals of *M. f. flavigula* feeding or searching for food (fruits, flowers, or insects). We also counted the times they were flycatching or feeding on human-provided food. As their diet in South America seems to be very different compared to the North and Central American populations (Stacey 1981, Kattan 1988, Sandoval 2016) - where its distribution is known to be very associated with the presence of granaries - we collected data on food habits and the usage of granaries.

#### Territory size and spatial analysis

To estimate territory sizes and foraging areas, we spent at least four days in each territory, and geo-referenced every point where one or several individuals were found to be foraging or static. We identified “hot-points” key-landmarks in each territory. After our initial geo-tagging, we quantified the frequency of visits to those “hot-points”. We also classified the points in the following categories: perch (PE) or foraging tree/structure (FO), granary (GR), cavity (CA), or a combination of any of the former structures. Finally, we recorded if the structures were natural (dead or living plants) or man-made (concrete/wooden utility post, concrete wall, or bird feeder).

To estimate territory size, we used both the Minimum Convex Polygon (MCP) and the Kernel density estimator (KDE) approaches using the R package “adehabitatHR” (Calenge 2006, Castaño et al. 2019). MCP consists on circumscribing all the outermost external location points collected (Mohr 1947), which gave us a rough but consistent estimation of the general area encompassed in each territory. While, KDE technique will estimate a frequency-based probability sketch of the internal use of the territory (Tingley et al. 2014), identifying core areas (areas of intense use) based on the utilization distributions generated by the package (Barg et al. 2004). As we used the “hot-points” method for mapping territories, our location points are clustered in specific areas or hot-points within the territories for which KDE can estimate core areas using the amount of location points registered in each “hot-point” but MCP will give a more conservative estimate of the total territory size.

## Results

### Group size and structure

The number of individuals per territory in our study area ranged from 5 to 9 individuals (7.5 ± 1.43 sd). The mean number of individuals in each sex category was 3.1 ± 1.10 for females, and 4.4 ± 0.96 for males. We found nine juveniles, and at least three chicks during our monitoring of territories. La Presidenta had one chick and two juveniles; La Frontera had one juvenile; Alejo had one chick and two juveniles; San Félix had one chick; CES had two juveniles; and Space had two juveniles. The number of adult individuals in each territory remained constant during our study (Table 1). In general, every group showed a cohesive pattern, especially at sunrise and sunset. Overall, we did not observe any beginning or strong power struggles (intense space competitions for dispersal purposes), neither in very close groups (i.e. Space and Baleares) nor between others (Figure 1). Nevertheless, more intense observations in each group should be done in order to give any result.

**Table 1.**
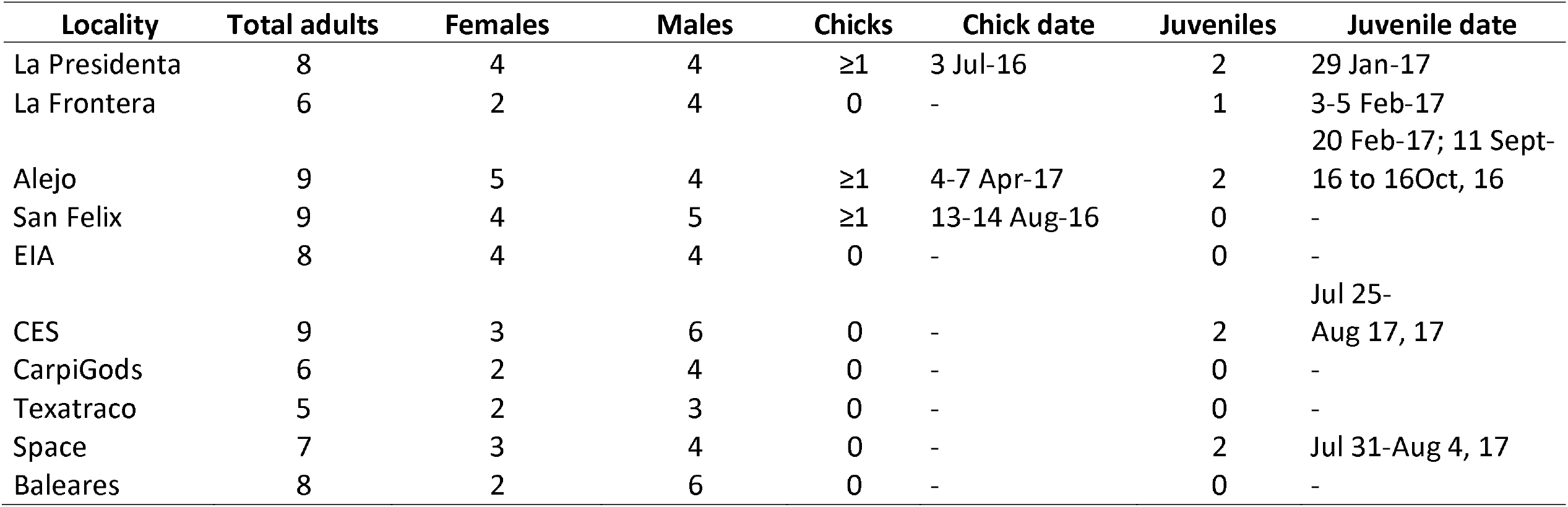
Group composition and group size of each studied territory. Estimated number of chicks and juveniles are shown along with their sighting dates.

### Tree species used for cavities and feeding

We found 53 species of trees (28 native and 25 exotic) representing 26 families and 428 individuals used by *M. f. flavigula* across our sampled locations. Ten species of trees and wooden utility posts were used for cavity-nesting, and 31 were used to forage and to feed. The other two categories include perches, which comprise a total of 44 species; and granaries, with 19 species (Table 2). In total we found 172 cavities, and in average 17.2 ± 8.2 cavities per group (Table 3). We recorded 539 feeding events, which include food items provided by tree species (fruit, insects in the bark/gleaning, or nectar from flowers, Table 4) and from behaviors like flycatching, and man-made food (i.e. rice, corn, or bread, Table 5). The diet of *M. f. flavigula* consisted primarily of insects from the bark (provided by 24 species of trees), being *A. carbonaria* the most visited tree; following by flycatching; flower visiting (provided by six species), being *Ochroma pyramidale* the most visited species; and finally, fruits (provided by 30 species), from which *Dypsis lutescens* was the preferred tree to visit (see Table 4).

**Table 2.** Trees and man-made structures used by *M. f. flavigula* in the Aburrá Valley. The more diverse family is Fabaceae (n = 9);, *A. carbonaria* was the tree species more often used for several daily activities such as a food provider (FO), cavities (CAV), granary (GRA) and perch (PE). This species was the most abundant and it was also present in most territories (7/10).

| Species | Family | Origin | FO | CAV | GRA | PE | n | Group |
| --- | --- | --- | --- | --- | --- | --- | --- | --- |
| *Acacia decurrens* Willd. | Fabaceae | exotic |  |  |  | x | 1 | 1 |
| *Albizia carbonaria* Britton | Fabaceae | native | x | x | x | x | 34 | 7 |
| *Alchornea* sp. | Euphorbiaceae | native |  | x | x |  | 7 | 1 |
| *Araucaria heterophylla* (Salisb.) Franco | Araucariaceae | exotic |  |  |  | x | 6 | 5 |
| *Archontophoenix cunninghamiana* (H.Wendl.) H.Wendl. & Drude | Arecaceae | exotic |  |  | x | x | 7 | 2 |
| *Bauhinia picta* (Kunth) DC. | Fabaceae | native | x |  |  | x | 6 | 2 |
| *Brugmansia × candida* Pers. | Solanaceae | exotic |  |  |  | x | 1 | 1 |
| *Caesalpinia pluviosa* DC. | Fabaceae | exotic | x |  |  |  | 1 | 1 |
| *Callistemon speciosus* (Sims) Sweet | Myrtaceae | exotic |  |  |  | x | 1 | 1 |
| *Calophyllum brasiliense* Cambess. | Calophyllaceae | native |  |  |  | x | 1 | 1 |
| *Cascabela thevetia* (L.) Lippold | Apocynaceae | native | x |  |  |  | 15 | 1 |
| *Cecropia angustifolia* Trécul | Urticaceae | native | x | x | x | x | 14 | 4 |
| *Cecropia telenitida* Cuatrec. | Urticaceae | native | x | x | x | x | 15 | 1 |
| *Citrus* sp. | Rutaceae | exotic | x |  |  | x | 1 | 1 |
| *Clusia* sp. | Clusiaceae | native | x |  |  |  | 1 | 1 |
| *Cordia* sp. | Boraginaceae | native |  |  |  | x | 2 | 1 |
| *Corymbia* sp. | Myrtaceae | exotic |  | x |  | x | 57 | 6 |
| *Croton magdalenensis* Müll.Arg. | Euphorbiaceae | native | x |  | x | x | 42 | 1 |
| *Cupressus lusitanica* Mill. | Cupressaceae | exotic |  |  |  | x | 8 | 3 |
| *Delostoma integrifolium* D.Don | Bignoniaceae | native |  |  |  | x | 1 | 1 |
| *Dry dead tree* | - | - |  | x | x | x | 3 | 2 |
| *Dyopsis lutescens* (H.Wendl.)<br>Beentje & J.Dransf. | Areaceae | exotic | x |  | x | x | 8 | 2 |
| *Electric wooden post* | - | - |  | x | x |  | 4 | 3 |
| *Erythrina fusca* Lour. | Fabaceae | native | x |  |  | x | 9 | 3 |
| *Erythrina poeppigiana* (Walp.)<br>O.F.Cook | Fabaceae | native |  |  | x | x | 2 | 2 |
| *Eucalyptus cinerea* F.Muell. ex<br>Benth. | Myrtaceae | exotic |  |  |  | x | 4 | 2 |
| *Eucalyptus* sp. | Myrtaceae | exotic |  | x | x | x | 2 | 2 |
| *Ficus* sp. | Moraceae | exotic | x |  |  | x | 2 | 2 |
| *Fraxinus uhdei* (Wenz.) Lingelsh. | Oleaceae | exotic |  |  | x | x | 14 | 2 |
| *Guzmania* sp. | Bromeliaceae | native | x |  |  |  | 7 | 2 |
| *Handroanthus chrysanthus*<br>(Jacq.) S.O.Grose | Bignoniaceae | native | x |  |  | x | 11 | 4 |
| *Inga* sp. | Fabaceae | native | x |  |  | x | 3 | 3 |
| *Jacaranda mimosifolia* D.Don | Bignoniaceae | exotic | x |  |  | x | 1 | 1 |
| *Leucaena leucocephala* (Lam.)<br>de Wit | Fabaceae | exotic | x |  |  | x | 4 | 1 |
| *Mangifera indica* L. | Anacardiaceae | exotic |  |  |  | x | 1 | 1 |
| *Quararibea cordata* (Bonpl.)<br>Vischer | Malvaceae | native | x |  |  | x | 1 | 1 |
| *Melaleuca quinquenervia* (Cav.)<br>S.T.Blake | Myrtaceae | exotic | x |  |  | x | 7 | 2 |
| *Meriania nobilis* Triana | Melastomataceae | native |  |  | x | x | 3 | 2 |
| *Musa* sp. | Musaceae | exotic | x |  |  |  | 4 | 1 |
| *Myrcia popayanensis* Hieron. | Myrtaceae | native | x | x | x | x | 12 | 2 |
| *Ochroma pyramidale* (Cav. ex Lam.) Urb | Malvaceae | native | x | x |  | x | 24 | 7 |
| *Pachira speciosa* Triana & Planch. | Malvaceae | native |  |  | x |  | 1 | 1 |
| *Passiflora mollissima* (Kunth) L.H.Bailey | Passifloraceae | native | x |  |  | x | 2 | 1 |
| *Persea americana* Mill. | Lauraceae | native | x |  | x | x | 7 | 3 |
| *Persea caerulea* (Ruiz & Pav.) Mez | Lauraceae | native | x |  | x | x | 20 | 8 |
| *Psidium guajava* L. | Myrtaceae | native | x |  |  | x | 8 | 2 |
| *Quercus humboldtii* Bonpl. | Fagaceae | native | x |  | x | x | 12 | 1 |
| *Schinus terebinthifolia* Raddi | Anacardiaceae | exotic |  |  |  | x | 6 | 1 |
| *Spathodea campanulata* P.Beauv. | Fabaceae | exotic | x | x | x | x | 12 | 3 |
| *Syagrus sancona* (Kunth) H.Karst. | Arecaceae | native | x |  |  |  | 5 | 1 |
| *Syzygium jambos* (L.) Alston | Myrtaceae | exotic |  |  |  | x | 1 | 1 |
| *Terminalia* sp. | Combretaceae | exotic |  |  |  | x | 1 | 1 |
| *Washingtonia robusta* H.Wendl. | Arecaceae | exotic | x | x | x |  | 3 | 2 |
| *Weinmannia* sp. | Cunoniaceae | native | x |  |  | x | 1 | 1 |
| *Zanthoxylum rhoifolium* Lam. | Rutaceae | native |  |  |  | x | 2 | 2 |

**Table 3.** Species of trees (and man-made structures) used for making cavities. The tree *Albizia carbonaria* was often used, with 18 of the 34 individuals hosting cavities.

| Species | Total individuals | Total used | Total cavities |
| --- | --- | --- | --- |
| *Albizia carbonaria* | 34 | 18 | 77 |
| *Alchornea* sp. | 7 | 1 | 1 |
| *Cecropia angustifolia* | 14 | 1 | 3 |
| *Cecropia telenitida* | 15 | 4 | 17 |
| *Corymbia* sp. | 57 | 3 | 7 |
| Dry dead tree | 3 | 2 | 5 |
| Wooden utility post | 4 | 3 | 10 |
| *Eucalyptus* sp. | 2 | 1 | 1 |
| *Myrcia popayanensis* | 12 | 2 | 10 |
| *Ochroma pyramidale* | 24 | 2 | 13 |
| *Spathodea*<br>*campanulata* | 12 | 3 | 6 |
| *Washingtonia robusta* | 3 | 1 | 22 |

**Table 4.** The use of different species of trees for feeding proposes varied extensively. Different species offer different feeding items/resources.

| Species | Fruit | Flower | Gleaning |
| --- | --- | --- | --- |
| *Albizia carbonaria* | 0 | 0 | 65 |
| *Bauhinia picta* | 0 | 0 | 3 |
| *Caesalpinia pluviosa* | 0 | 0 | 8 |
| *Cascabela thevetia* | 0 | 0 | 8 |
| *Cecropia angustifolia* | 0 | 0 | 13 |
| *Cecropia telenitida* | 0 | 0 | 4 |
| *Citrus* sp. | 0 | 0 | 1 |
| *Clusia* sp. | 0 | 0 | 4 |
| *Croton magdalenensis* | 0 | 0 | 4 |
| *Dypsis lutescens* | 12 | 0 | 0 |
| *Erythrina fusca* | 8 | 5 | 5 |
| *Ficus* sp. | 0 | 0 | 3 |
| *Guzmania* sp. | 0 | 8 | 0 |
| *Handroanthus chrysanthus* | 0 | 0 | 20 |
| *Inga* sp. | 0 | 0 | 1 |
| *Jacaranda mimosifolia* | 0 | 0 | 3 |
| *Leucaena leucocephala* | 0 | 0 | 10 |
| *Quararibea cordata* | 0 | 0 | 3 |
| *Melaleuca quinquenervia* | 0 | 0 | 2 |
| *Musa* sp. | 0 | 3 | 0 |
| *Myrcia popayanensis* | 0 | 0 | 4 |
| *Ochroma pyramidale* | 0 | 31 | 11 |
| *Passiflora mollissima* | 0 | 7 | 0 |
| *Persea americana* | 0 | 0 | 14 |
| *Persea caerulea* | 0 | 0 | 26 |
| *Psidium guajava* | 2 | 0 | 0 |
| *Quercus humboldtii* | 6 | 0 | 0 |
| *Spathodea campanulata* | 0 | 9 | 5 |
| *Syagrus sancona* | 0 | 0 | 1 |
| *Washingtonia robusta* | 6 | 0 | 1 |
| **Total** | **28** | **63** | **218** |

**Table 5.** Feeding preferences by groups of *M. f. flavigula* in the Aburrá Valley. From the 539 feeding sightings, gleaning in trees (n = 214) was the most common behavior, following of man-provided food (n =125).

| Group | Fruit | Man-provided | Gleaning | Flower | Flycatching |
| --- | --- | --- | --- | --- | --- |
| La Presidenta | 5 | 72 | 15 | 3 | 27 |
| La Frontera | 0 | 0 | 12 | 5 | 8 |
| Alejo | 4 | 9 | 64 | 12 | 5 |
| San Félix | 1 | 8 | 15 | 12 | 25 |
| EIA | 6 | 24 | 11 | 0 | 3 |
| Universidad CES | 7 | 0 | 28 | 14 | 11 |
| CarpíGods | 0 | 9 | 16 | 7 | 12 |
| Texatraco | 0 | 0 | 20 | 4 | 0 |
| Space | 0 | 3 | 16 | 3 | 10 |
| Baleares | 13 | 0 | 17 | 3 | 0 |

### Tree species used as granaries

In the Aburrá Valley, *M. f. flavigula* uses several species of trees to construct their granaries. We found granaries in 19 species of trees but also in man-made structures such as electric posts (Table 1). We found that out of 42 granaries, 32 were actively used, and 34 were living trees (Table 6). We found granaries in nine out of 10 localities, in both urban and rural areas. La Frontera was the only group where we did not find any granaries, and San Félix was the group that held more granaries (n=10) (Figure 1). We found that *A. carbonaria* was the most used tree, with seven (out of 42) trees used as granaries, and distributed in seven of the 10 groups: where four were actively used, and six were living trees. We also found two trees we couldn’t identify used as granaries, from which only one was active. Likewise, we found one active wooden utility post and crevices of a brick-building used for food-storing.

**Table 6.** Trees species and other structures used as granaries. *A. carbonaria* prevailed as the most frequently use tree for granary-construction.

| Species used as granaries | Used (n) | Active/not active | Alive/dead | Locations |
| --- | --- | --- | --- | --- |
| *Albizia carbonaria* | 7 | 4/3 | 6/1 | 7 |
| *Alchornea* sp. | 2 | 2/- | 2/- | 1 |
| *Archontophoenix cunninghamiana* | 1 | 1/- | 1/- | 2 |
| *Cecropia angustifolia* | 2 | 2/- | 1/1 | 4 |
| *Cecropia telenitida* | 4 | 4/- | 4/- | 1 |
| *Croton magdalenensis* | 1 | 1/- | 1/- | 1 |
| *Dry dead tree* | 2 | 1/1 | -/1 | 2 |
| *Dyopsis lutescens* | 1 | 1/- | -/1 | 2 |
| *Wooden utility post* | 1 | 1/- | -/1 | 3 |
| *Erythrina poeppigiana* | 1 | 0/- | 1/- | 2 |
| *Eucalyptus sp.* | 2 | 1/1 | -/2 | 2 |
| *Fraxinus uhdei* | 1 | 1/- | 1/- | 2 |
| *Meriania nobilis* | 1 | -/1 | 1/- | 2 |
| *Myrcia popayanensis* | 3 | 3/- | 3/- | 2 |
| *Pachira speciosa* | 1 | 1/- | 1/- | 1 |
| *Persea americana* | 3 | 2/1 | 3/- | 3 |
| *Persea caerulea* | 3 | 1/2 | 3/1 | 8 |
| *Quercus humboldtii* | 2 | 2/- | 2/- | 1 |
| *Spathodea campanulata* | 2 | 2/- | 2/- | 3 |
| *Washingtonia robusta* | 2 | 2/- | 1/1 | 2 |

### Territory sizes

We found that territory sizes varied across the 10 studied groups According to the MCP estimated areas, some territories (i.e. Alejo) were at least five times bigger than others (La Presidenta, Figure 3 and Figure 4). The territories in rural areas (EIA) were bigger than the ones in strongly urbanized areas (CES), where some groups are very close to each other (< 1 km between main cavity/granary). In urban territories, birds moved freely across empty spaces between buildings, city parks and isolated trees. In rural areas, birds move between “hot-points” predictably with no trend to explore new foraging or perching trees. In both, urban and rural areas, birds constantly returned to a tree with cavities or to a granary (despite not being banded individuals).

**Figure 2.**
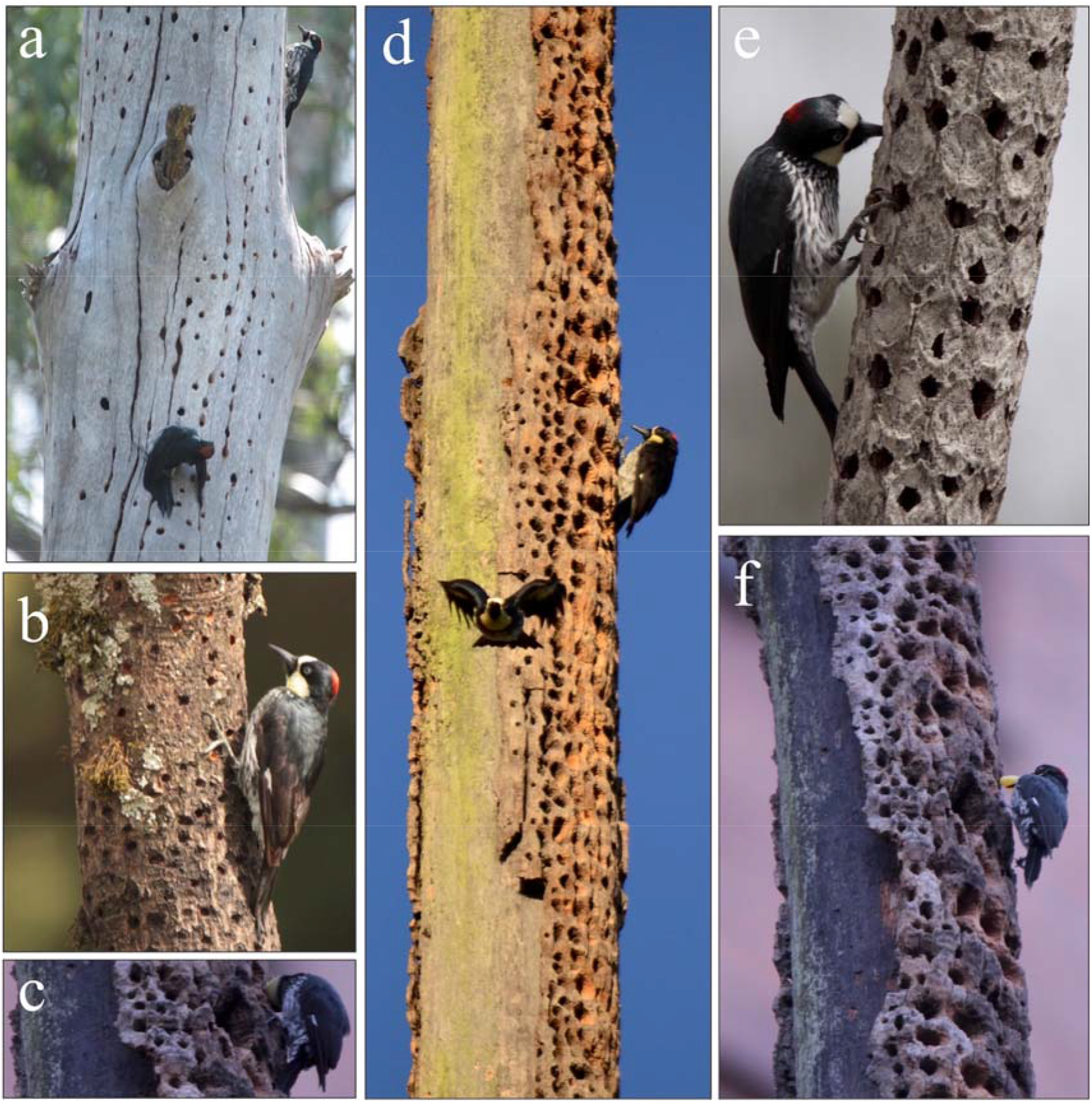
a-f. Active granaries used by *Melanerpes formicivorus* in three different groups. In *Eucalyptus* sp. (a); *Myrcia popayanensis* (b); *Washingtonia robusta* (c,d,f); and *Carica goudotiana* (e); Inds. Storing man-provided food (c,f). Photos: HLO & Paula Saravia.

**Figure 3.**
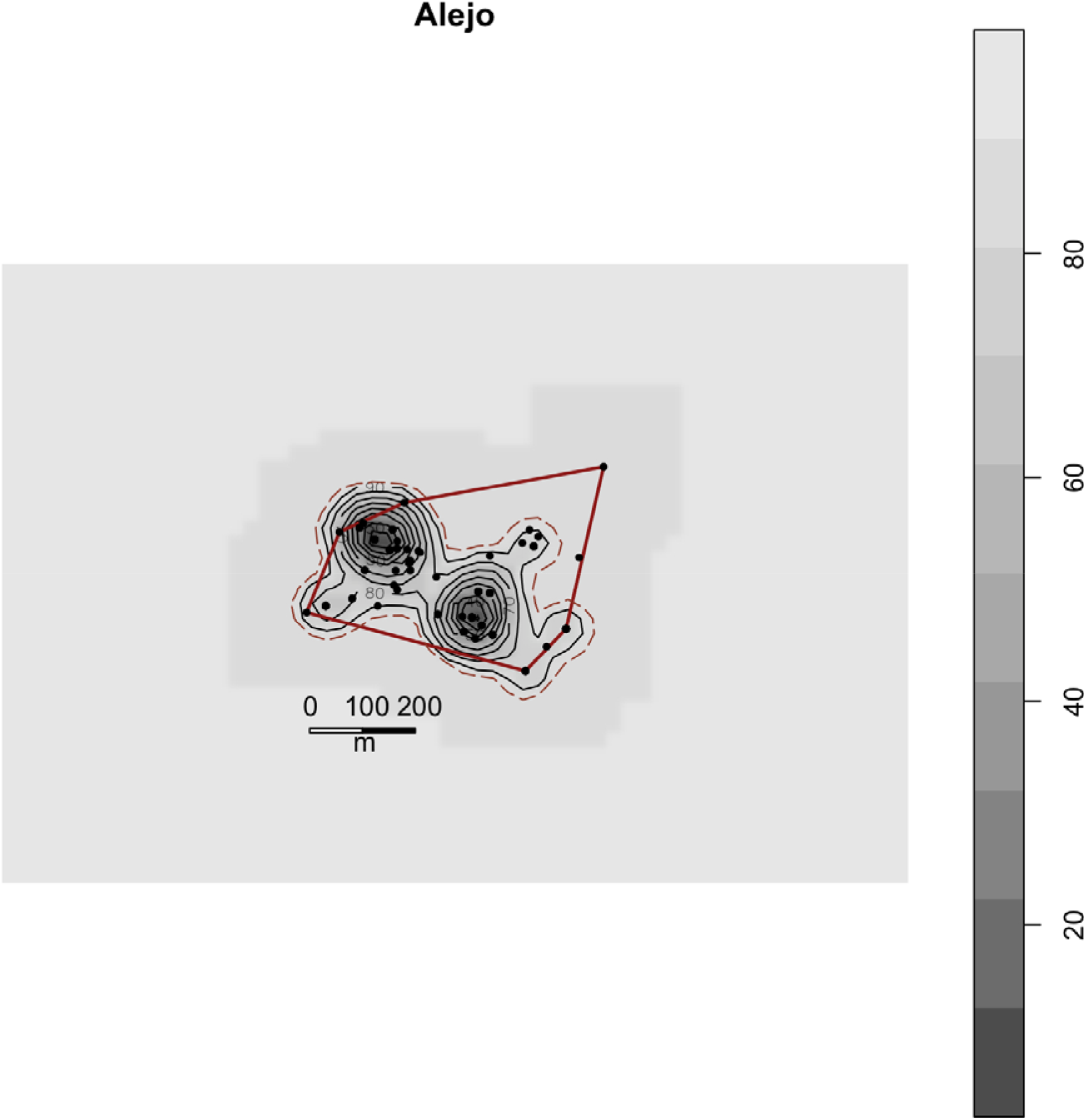
Estimated MCP and Kernel areas of movement/foraging for the groups Alejo and La Presidenta. External polygon represents the MCP, while gray shaded areas the Kernel distributions. The locations where the woodpeckers were registered and located several times are represented with the black dots.

**Figure 4.**
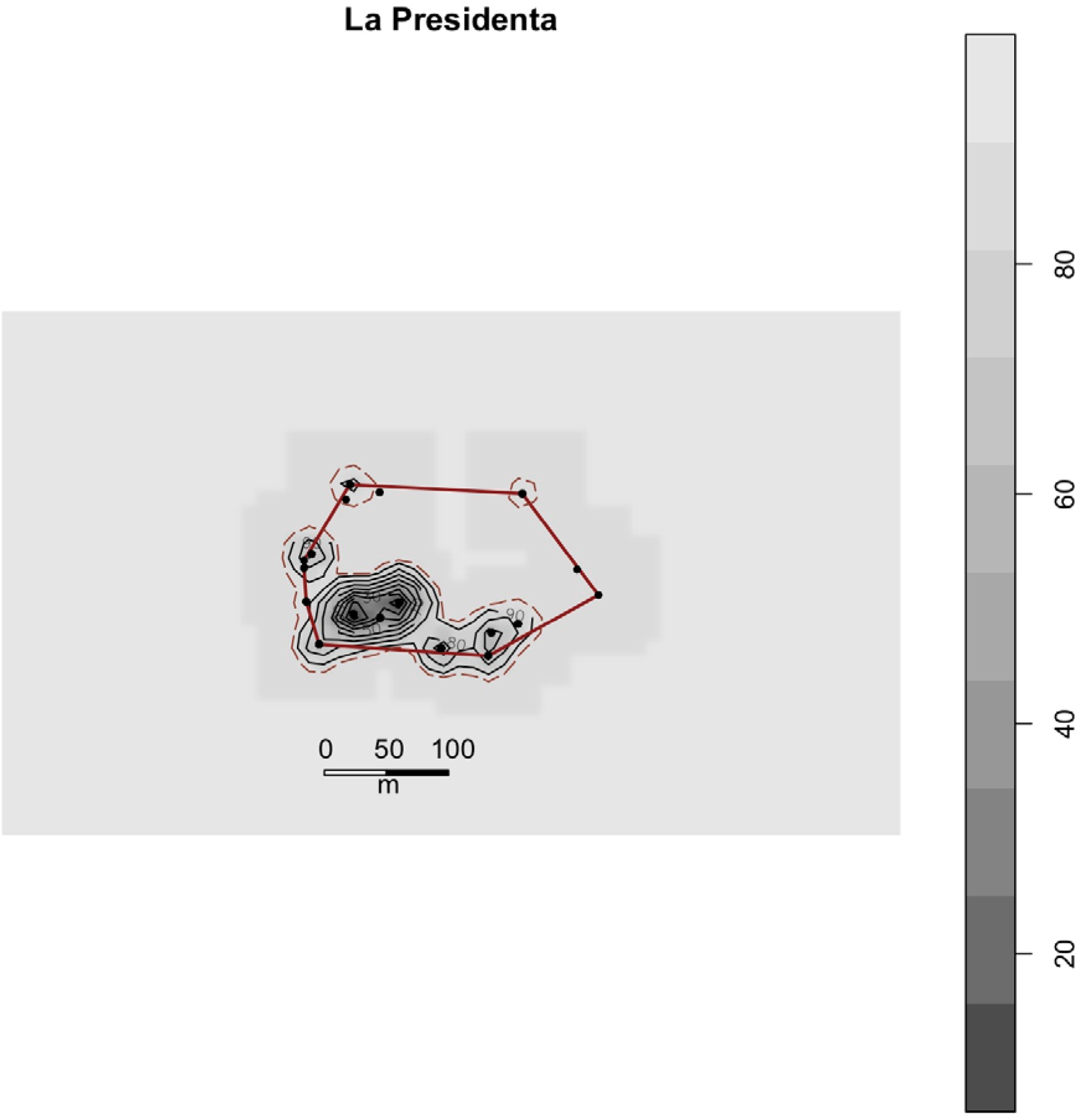
Estimated MCP and Kernel areas of movement/foraging for the groups Alejo and La Presidenta. External polygon represents the MCP, while gray shaded areas the Kernel distributions. The locations where the woodpeckers were registered and located several times are represented with the black dots.

## Discussion

Widely distributed bird species across latitudinal, altitudinal or habitat gradients are expected to be exposed and to explore different habitats and resources (Martin & Fitzgerald 2005). Therefore, a generalist strategy seems to be more suitable than specialization (either on resources or habitat) for such species (Sol et al. 2002). Our main goal was to characterize the habitats used by the Acorn Woodpecker (*Melanerpes formicivorus flavigula*) in urban and rural areas in northern South America. We found support for the hypothesis that *Melanerpes f. flavigula* is not limited by the presence of the Colombian Oak, *Quercus humboldtii* in urbanized or rural areas, at least in the Aburrá Valley region, where this species thrives in highly modified areas. Instead, our results suggest the Acorn Woodpeckers use a wide range of native and exotic tree species, particularly *Albizia carbonaria* (family: Fabaceae) that provides food, and surfaces for the construction of cavites and granaries or for perching.

We found this tree species near or inside areas occupied by Acorn Woodpeckers in the east slope of the Aburrá Valley. The use of a wide range of tree species might be related to the higher diversity of plants, therefore there may be a greater number of exploitable tree resources in Colombia, in comparison to other plant communities where the Acorn Woodpecker is also distributed (Mutke & Barthlott 2007). The use of other tree species might also be associated with the reduction of oak diversity: while in North America the Acorn Woodpecker exploits at least eight species of oaks (MacRoberts & MacRoberts 1976, Hooge et al. 1999, Johnson & Rosenberg 2006, Scofield et al. 2011), and in Central America it uses at least nine species (Skutch 1969, Stacey 1981, Stanback 1989, Rosas et al. 2008), in South America there is only one species of oak, *Quercus humboldtii* (Kattan 1988). Theferore, our study presents a wider perspective on the use of habitat and tree resourses by *M. formicivorus* in a highly urbanized area, where a mixture of native and exotic trees coexist, and where oaks are not abundant.

Consequently, our results support our second hypothesis: for the first time we provide evidence on use and construction of granaries in Colombia, specifically in the Aburrá Valley region. Although Miller (1963) found acorn stores and old nest holes in a *Cecropia* sp. tree, Kattan (1988) did not find any granaries and proposed that the construction of these granaries was unnecessary. Both studies were conducted in the southernmost distribution of this species in Colombia; our observation of the use of granaries gives reason to rethink several aspects about this species, including its natural history, distribution, and ecology. Local groups of Acorn Woodpeckers in the Aburrá Valley have retained this behavior, which was originally thought to be present only in temperate regions (MacRoberts & MacRoberts 1976, Koenig & Walters 2014), despite the apparent lack of seasonal changes that favors the storing of food for the winter months in North America (Koenig & Mumme 1987). Acorn Woodpeckers use both natural (*A. carbonaria* as the species most used as a granary) and artificial structures (like utility posts and brick buildings) for granaries in the Aburrá Valley in urban, peri-urban, and rural environments; which suggests this behavior of using granaries to store food (MacRoberts & MacRoberts 1976), and thus shape the group structures, to enhance reproduction (Koenig & Mumme 1987), and to predict the distribution of this species at local scales (Johnson & Rosenberg 2006) is widespread across the Americas.

One limitation of this study in relation with the usage of granaries is the lack of information on food storage. Because we did not analyze which foods were stored in the granaries, we were not able to evaluate whether food storage in granaries limits the distribution of this species. We found, however, that granaries are not saturated (at least not visually) with food throughout the year, suggesting that the main role of the granaries is related to social cohesion of family groups.

We found behavioral similarities among our study groups and the ones studied in North America; for example, our studied groups tended to construct granaries in trees with small trunks (palm trees in this case) such as in *Washingtonia robusta* H.Wendl., *Dypsis lutescens* (H.Wendl.) Beentje & J.Dransf., and *Archontophoenix cunninghamiana* (H.Wendl.) H.Wendl. & Drude, and also in dry secondary branches of other trees. This flexibility of making granaries in different sizes, parts and species of trees, drives us to conclude this species does not rely on Q. *humboldtii* in urbanized and rural areas in northern Colombia, as previously suggested (Freeman & Mason 2015), and that it uses a wider diversity of trees to make granaries, apart of feeding, nesting and foraging.

Moreover, since the diet of *M. formicivorus* consisted primarly of insects extracted from the bark of trees, and *Albizia carbonaria* was the most visited tree for this feeding category, we believe this might be an important factor in explainig their strong association. Further studies need to identify and quantify the insects present in this species of tree, and particularly to understand which ones made part of the diet of the Acorn Woodpecker. Whether *A. carbonaria* have any morphological and phenological trait that attracts insects is yet to be discovered, however, we found insects in different stages that can live in their irregular and rough bark, and pollinators and insect visitors keep close in their long flowering period from March to November (UICN-ORMAACC 2015). *Albizia carbonaria* also offers resources for cavity construction; it is usual to find living adult trees full of dry, sturdy branches (Varón & Morales 2016), which were abundant in our study area and are also a preferred characteristic by Acorn Woodpeckers when looking to start a cavity hole (MacRoberts & MacRoberts 1976).

The diet of *M. formicivorus* was variable, but there was a preference for insects, which seems to be a common trait through its distributional range (Winkler & Christie 2019). However, in temperate regions this species changes its diet to the consumption of acorns of various species of oaks when the acorn season begins, which generally starts in autumn (Koenig & Mumme 1987), when temperatures start declining, along with the availability of insects. Since there is no such climatic seasonality in Colombia (e.g. cold winters and hot summers), we believe there is no lack of insects throughout the year and instead, these populations could be also benefited by other resources like fruits, flowers, and man-provided food. This last kind of food (rice, bread, corn, dog food, bananas or even Cheetos) was a common behaviour in our study. Although this needs to be studied more rigurously, we believe it is an important resource at least in our sampled groups, both in urban or rural areas. For example, chicks of two groups (San Félix and La Presidenta) were being fed with this type of food.

The visiting of flowers is another feeding preference. In 1988, Kattan reports *M. formicivorus* visiting flowers of *Spathodea campanulata*, along with *Ochroma pyramidale* trees; the latter tree species as a preference was also documented during our observations. Both of this flowers have wide and big petals that seems to be attractive for this birds either to consume its nectar, and/or the water saved after a rain. The last types of food include fruits and seeds, which are specially from: the exotic palm trees *Washingtonia robusta* and *Dypsis lutescens*, that are commonly planted in cities as an ornamental tree (Varón & Morales 2016), and also from Q. *humboldtii* oaks, from which we saw Acorn Woodpeckers collecting acorns in the only group that had oaks within their territory. Morover, although Kattan (1988) reported the consumption of sap in oak trees by *M. formicivorus* in south west of Colombia, we never saw this behavior during our study, neither in oaks or other trees. However, locals from the group San Félix told us they have seen this woodpecker “entering their tongues like sucking the tree’s sap” (referring to *Persea americana* and *Myrcia popayanensis* trees) in holes that are smaller than granary ones. Finally, although consuming acorns from Q. *humboldtii* is a known behavior in Colombia (Kattan 1988), this is the first time reporting a usage of oaks for granary construction.

For the first time, we describe the use and construction of granaries in Colombia, specifically in the Aburrá Valley region, for which *M. formicivorus* uses a wide variety of tree species, and other alternatives like utility posts, building bricks and dead trees in all types of environments, urban, peri-urban, and rural areas. This behavior was originally thought to be present only in temperate regions (MacRoberts & MacRoberts 1976, Koenig & Walters 2014), but we believe this could be a common yet not studied behavior in the Colombian Andes. We believe granaries play an important role in this woodpecker’s social structure and establishment since our studied populations didn’t make as much holes or stored as much food as they do in northern populations (Koenig & Mumme 1987), but they did defend the granaries from other individuals or species and frequently visited them and gathered. Finally, although nesting was not studied, it is likely that the birds breed communally in Colombia as elsewhere in their range.

In conclusion, our study indicates that the Acorn Woodpecker is succesfully ocupying urban areas in tropical latitudes where it reproduces, make granaries and feeds on a wide variety of resources. We believe that availability of tree species, specially *A. carbonaria* that offers food (insects, flowers and fruits), home (cavities), and granaries is key to explain the presence of this species in this highly urbanized area. There is a need for more studies in both slopes of the Aburrá Valley and nearby areas surroundings, where basic natural history and ecology are poorly known, and which might provide a better understanding of the drivers and limitations of the distribution of this species in a wider regional scale. Particularly, in locations where *Q. humboldtii* trees are present might be key to understand the dependence of usage of this oak in the Northern Andes.

## ACKNOWLEDGMENTS

We specially thank Paula Saravia, Catalina Arenas, and other volunteer students from the Facultad de Ciencias y Biotecnología from Universidad CES that provided logistical support for this study. We thank Paula Saravia for allowing us using her pictures. We are also in debt with Prof. Dino Tuberquia and student Lina Bolívar for kindly helping us identify all the species of trees. María Castaño assisted with the estimation of MCP and Kernel. Finally, to the owners of more than 20 private areas that generously allowed us to work on their land.

